# Development of a physiological insulin resistance model in human stem cell-derived adipocytes

**DOI:** 10.1101/2022.02.22.481495

**Authors:** Max Friesen, Andrew S. Khalil, M. Inmaculada Barrasa, Jacob F. Jeppesen, David J. Mooney, Rudolf Jaenisch

## Abstract

Adipocytes are key regulatory cells of human metabolism, and their dysfunction in insulin signaling is central to metabolic diseases such as type II diabetes mellitus (T2D). However, the progression of insulin resistance that leads to T2D is still poorly understood. This limited understanding is due, in part, to the dearth of suitable models of insulin signaling in human adipocytes. Traditionally, in vitro adipocyte models fail to recapitulate in vivo insulin signaling, possibly due to exposure to supraphysiological nutrient and hormone conditions. Here, we have developed a sensitization protocol for human pluripotent stem cell-derived adipocytes that uses physiologically relevant nutrient conditions to produce a potent signaling response comparable to in vivo adipocytes. After systematically optimizing conditions, this protocol allows for robust insulin-stimulated glucose uptake and transcriptional insulin response. Furthermore, exposure of these sensitized adipocytes to physiologically relevant hyperinsulinemic conditions dampens insulin-stimulated glucose uptake and dysregulates transcription of insulin-responsive genes. Overall, this sensitization methodology provides a novel platform for the mechanistic study of insulin signaling and resistance using human pluripotent stem cell-derived adipocytes.

**Teaser:** A new protocol to generate hPSC-adipocytes that respond to physiological insulin levels and can model diabetes.

## Introduction

The increasing prevalence of metabolic diseases(*1, 2*) has lead to a commensurate interest in mechanistically understanding how environment and nutrition affect metabolism. This is due to evidence that these factors have been shown to regulate human metabolism and contribute to metabolic dysfunction, including insulin resistance and diabetes(*3, 4*). However, despite the significance of metabolic diseases in global public health, the tools for mechanistically studying the role of non-genetic factors in human metabolic dysfunction remain limited and insufficient. For example, current tools either rely on murine models or human in vitro cell culture systems, both of which poorly mimic in vivo human metabolism(*5*). A key limitation of in vitro cell culture systems is that the nutrient ingredients in culture media do not recapitulate in vivo human tissues(*6*). Despite multiple previous attempts to approximate human plasma with improved cell culture media, current in vitro culture models often still include specific nutrients at supraphysiological levels or have not yet been validated for insulin signaling response(*7*).

Adipose tissue is central to metabolic health and disease(*8*). The dysregulation of adipocyte function and loss of insulin sensitivity often observed in obesity is one of the key risk factors for the development of T2D(*9*). However, in vitro adipocyte culture models poorly mimic the functional capabilities of in vivo adipose tissue for the study of insulin resistance. Most notably, in vitro human adipocyte cultures lack the capacity to perform insulin-stimulated glucose uptake at sufficient levels to observe how environmental or nutritional factors might influence this process. Insulin-mediated glucose uptake and transcriptional response is primarily effectuated by AKT2 after its phosphorylation downstream of the insulin receptor(*10*). Previous in vitro models have generally been able to show reasonable AKT2 phosphorylation induced by insulin, but this does not translate into physiologically relevant insulin-stimulated glucose uptake. Overall, these insufficiencies of in vitro culture models limit the ability to mechanistically study the dysregulation and loss of insulin sensitivity in human adipocytes.

Human pluripotent stem cells (hPSCs) are a powerful tool for pathobiology due to their ability to differentiate into almost any tissue to model various diseases, the ease of implementation of genome editing tools, and exquisite control over their culture conditions(*11*). They also have unlimited availability, which is a major advantage over primary tissues that can be notoriously hard to obtain. This previous limitation is especially true in the case of metabolically healthy adipose tissue. Previously, an established adipocyte differentiation protocol(*12*) recapitulated several key adipocyte phenotypes in terms of gene expression and lipid accumulation. While these adipocytes have been used to study obesity and their contribution to cardiovascular disease phenotypes(*13, 14*), they poorly recapitulate insulin signaling. Specifically, insulin-stimulated glucose uptake in hPSC-derived adipocytes has been previously performed at supraphysiological maximal insulin stimulation and portrayed after blocking and subtraction of basal glucose uptake, artificially inflating the observed fold changes. The lack of insulin response at physiologically relevant levels is a key hurdle to overcome to fully unlock the potential of hPSC-derived adipocytes in metabolic disease modeling.

Here, we hypothesized that supraphysiological concentrations of key nutrients in cell culture medium, especially glucose and insulin, negatively impact the function of in vitro adipocytes.

Using an established hPSC adipocyte differentiation protocol, we employed a Design of Experiments (DoE) approach to identify optimal nutrient conditions to potentiate insulin-stimulated AKT2 phosphorylation and glucose uptake in hPSC-adipocytes. This optimized sensitization protocol resulted in a physiologically relevant culture condition that improved the functional maturation of the hPSC-derived adipocytes. Adipocytes produced via this approach also demonstrated robust transcriptional response upon insulin stimulation in multiple relevant metabolic pathways. Furthermore, we generated hyperinsulinemia-induced insulin resistance in adipocytes by modulating the insulin concentration during the final differentiation step. These insulin-resistant adipocytes demonstrated impaired insulin-stimulated glucose uptake and transcriptional responses. Our physiologically relevant adipocyte culture model provides a new potential platform for mechanistic studies into how environmental and nutritional factors influence insulin signaling and resistance in human adipocytes.

## Results

### Protocol optimization to generate insulin-responsive adipocytes

To enhance the insulin response of hPSC-adipocytes, we sought to optimize an established protocol(*12*) by adding a sensitization step with optimized nutrient and hormone conditions and length of culture (Fig. 1a). We started with a serum-free DMEM-base medium and sought to determine the ideal concentrations of glucose, insulin, albumin, and IGF to maximize insulin-stimulated glucose uptake and AKT2 phosphorylation. We employed a DoE approach to explore the relationship between these factors systematically and generated 26 different medium compositions (Supplementary Table 1), with which we treated adipocytes for one week. We compared these conditions to the results we obtained using the published differentiation medium in both AKT2 phosphorylation (Fig. 1b) and glucose uptake (Fig. 1c) and modeled the nutrient interactions to determine the principal components influencing the insulin responses and their desired concentrations. Our model indicated that the principal parameters driving both the AKT2 phosphorylation (Fig. 1d) and glucose uptake (Fig. 1e) are insulin and glucose, with IGF and albumin having negligible effects (Supplementary Fig. 1).

**Fig. 1:**
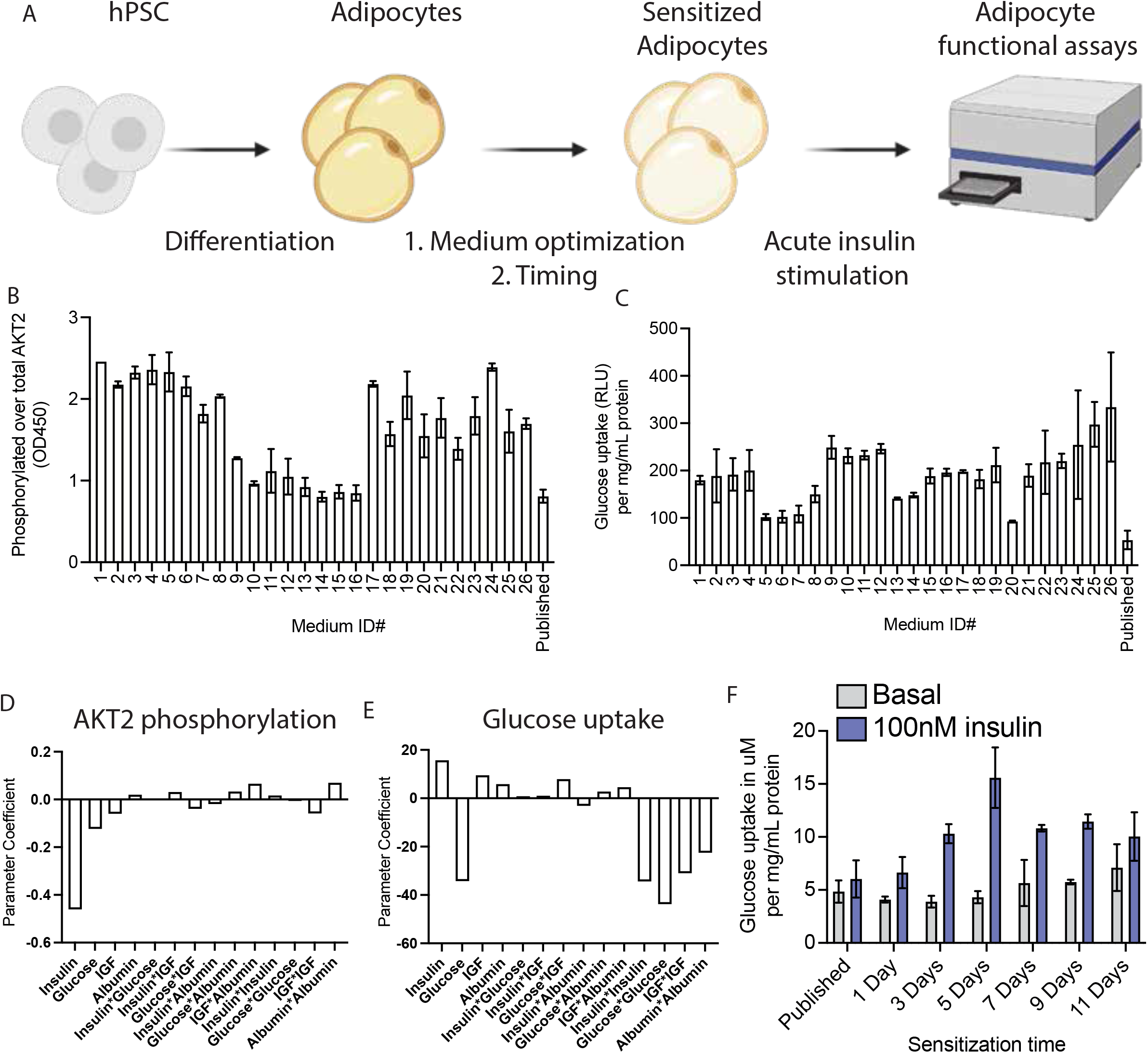
Generation of an insulin-sensitive human adipocyte model. A, Schematic indicating the experimental setup to potentiate insulin response in functional assays. hPSCs are differentiated into adipocytes using a published protocol, after which an additional step was added and optimized to sensitize adipocytes. B-C, Phosphorylation of AKT2 normalized to total AKT2 (B) and glucose uptake (C) after 100nM insulin stimulation measured for all the medium compositions from our DoE model and the medium from a previously published protocol. D-E, The parameter coefficient of each factor in our DoE model indicating its contribution to the AKT2 phosphorylation (D) and glucose uptake (E). F, Timecourse of sensitization with the DoE-optimized media measuring glucose uptake at baseline and after insulin stimulation. Results are normalized to total protein for each sample. All bar graphs depict the mean with error bars representing s.d., n=2 biological replicates.

Informed by the DoE model, we generated a sensitization medium containing 1g/L glucose and 100pM insulin. Notably, these concentrations were in the range of what one would find in the plasma of a healthy human(*15*). We then optimized the exposure time to the sensitization medium, with a clear maximum in insulin-stimulated glucose uptake reached after five days of sensitization, with minimal effects on basal glucose uptake (Fig. 1f). This sensitization protocol also resulted in potent AKT2 phosphorylation (Supplementary Fig. 1). Published literature frequently uses overnight serum starvation, which could roughly be compared to our one-day sensitization time, indicating a present but weak insulin response in line with previous studies. No condition in the DoE screen or the exposure duration time points resulted in observable adverse effects on the adipocytes as measured by total protein or total AKT2. The finalized sensitization protocol also did not reduce adipocyte numbers or change their accumulated lipid content (Supplementary Fig. 1). With this optimized media and five-day sensitization protocol, the hPSC-adipocytes displayed a >3-fold increase in glucose uptake upon insulin stimulation.

### Optimized medium permits adipocyte response to physiological insulin levels

As shown in figure 1, the sensitized adipocytes responded strongly to maximal stimulation with 100nM insulin, a standard concentration used in many previous publications. However, given our sensitization medium’s physiologically relevant baseline insulin concentration, we sought to identify whether insulin levels resembling human postprandial serum concentrations could also lead to insulin response in sensitized adipocytes (Fig. 2a). In an insulin dose-response assay, we demonstrated robust AKT2 phosphorylation at single-digit nanomolar concentrations of insulin in our sensitized adipocytes, with negligible response using previously published media (Fig. 2b). At a postprandial physiologically relevant stimulation of 10nM insulin for ten minutes(*16*), we observed translocation of GLUT4 from the intracellular space to the cell membrane (Fig. 2c). Relative to baseline membrane GLUT4, we measured a 71% increase in membrane-resident signal via total internal reflection microscopy (TIRF) imaging (Fig. 2d).

**Fig. 2:**
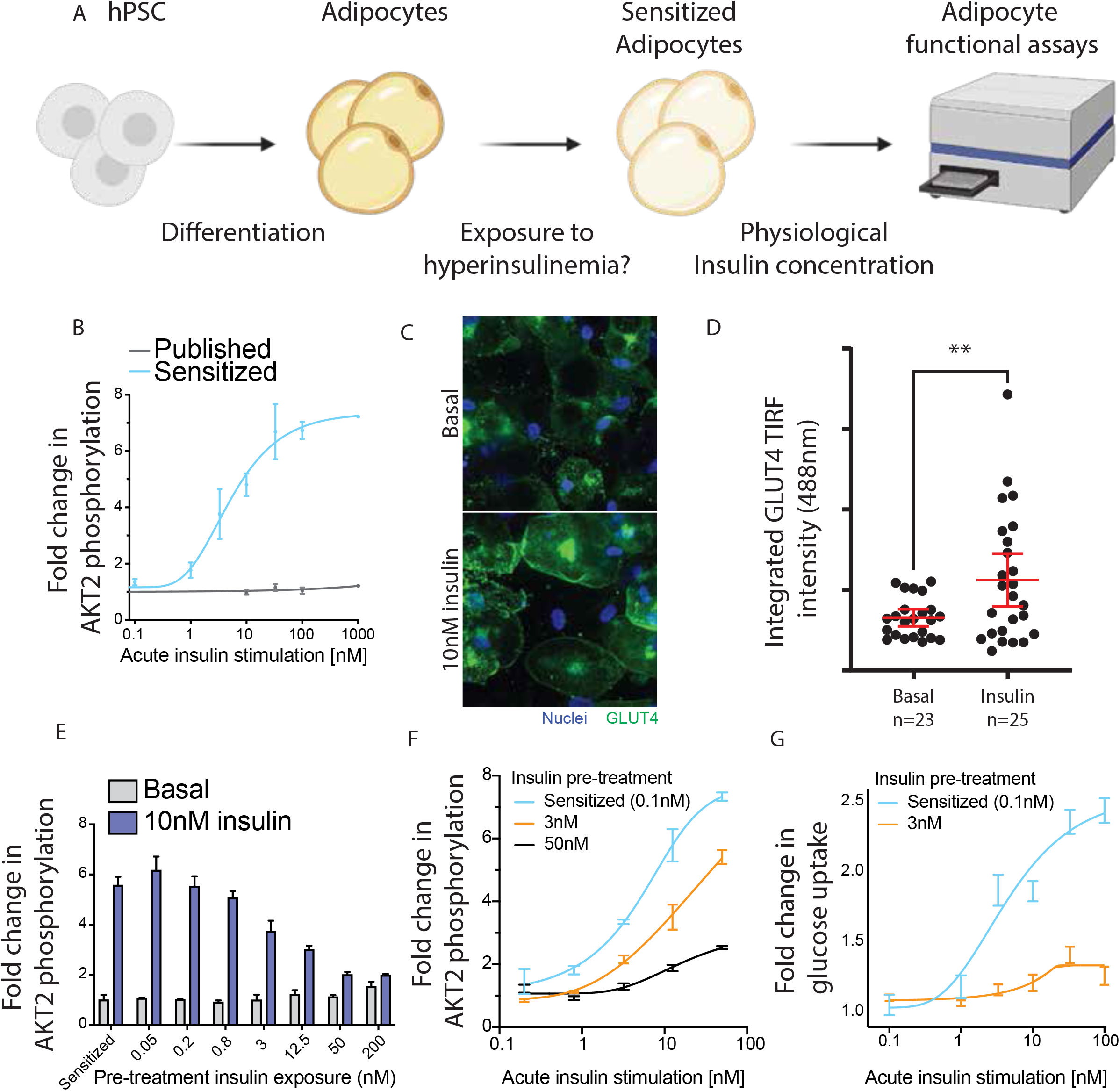
Physiological insulin levels induce insulin response and resistance. A, Schematic indicating the experimental setup to measure insulin dose-response and induction of insulin resistance. B, Insulin dose-response curve showing fold change in AKT2 phosphorylation compared to the unstimulated state. C, Representative images of GLUT4 translocation to the cell membrane upon insulin stimulation. D, TIRF measurement of GLUT4 signal intensity at the adipocyte cell membrane (**=p<0.01, unpaired two-tailed T-test). E, AKT2 phosphorylation in basal and insulin-stimulated state after increasing exposure to higher insulin levels during the sensitization period. Results are normalized to total AKT2 and plotted as fold change compared to the sensitized basal state. F, Insulin dose-response curve showing AKT2 phosphorylation fold change in three insulin pre-exposure conditions. Results are normalized to total AKT2 and plotted as fold change to unstimulated cells in that condition. G, Insulin dose-response curve showing glucose uptake for sensitized and hyperinsulinemia-exposed adipocytes. Results are normalized to total protein content and plotted as fold change to unstimulated cells. Bar graphs depict the mean with error bars representing s.d., dose-response curves depict a nonlinear fit curve with error bars representing s.d., scatterplot depicts individual cell values with mean and 95% CI overlaid, n=3 biological replicates unless otherwise indicated.

### Hyperinsulinemia induces insulin resistance in adipocytes

After validating normal insulin responsiveness in the sensitized adipocytes, we sought to establish a model of hyperinsulinemia-induced insulin resistance. To induce this insulin resistance, we exposed adipocytes throughout the sensitization period to various insulin concentrations. This exposure significantly dampened phosphorylation of AKT2 commensurate to the insulin level (Fig. 2e). AKT2 phosphorylation fold changes were decreased across the entire insulin dose-response curve when exposed to a physiologically relevant chronic level of 3nM insulin, mimicking systemic hyperinsulinemia in T2D patients (Fig. 2f). At supraphysiological maximal (100nM) insulin stimulation, the response was reduced by approximately 15%, while at 0.8nM and 3.2nM acute insulin stimulation AKT2 phosporylation was decreased by about 50-80%. Exposure to 3nM of insulin for the final five days of differentiation almost completely abrogated any insulin-responsive glucose uptake (Fig. 2g). The degree of induced insulin resistance correlated with the concentration of insulin used during the sensitization period and was reproducible in one additional ESC and two iPSC lines (Supplementary fig. 2). In summary, these results demonstrate that the sensitized adipocytes are insulin sensitive with an appropriate dose-response to physiologically relevant nanomolar exposures and that this sensitivity can be blunted by exposure to hyperinsulinemia, leading to insulin resistance both in signaling and function.

### Transcriptomic characterization of insulin-responsive and -resistant adipocytes

Having validated functional insulin responses in the sensitized adipocytes, we examined the transcriptional changes induced by insulin stimulation. We first verified the transcriptional adipocyte identity of the sensitized protocol developed here in comparison to adipocytes differentiated using a published protocol(*12*) and to primary adipocytes isolated from non-obese, non-diabetic humans. On a global transcriptomic level, the sensitized hPSC-derived adipocytes were equivalent to cells derived using published protocols in their adipocyte identity, with similar principal component distances observed relative to the primary adipocytes (Supplementary fig. 3). There was also strong concordance between all samples in their expression of core adipocyte genes as identified by the Human Protein Atlas(*17*). Despite similar transcriptomic adipocyte identities during basal culture, we explored how transcriptional responses after insulin stimulation might differ between adipocytes derived using published methods and the cells described here. We generated three groups of adipocytes; using the published protocol, as well as the sensitized (0.1nM), and chronic hyperinsulinemia (3nM) insulin-resistant cells described above. We observed that adipocytes derived using a standard published protocol show little to no transcriptional response upon insulin stimulation, while both the sensitized and insulin-resistant adipocytes responded potently to stimulation with 10nM of insulin for 4 hours (Supplementary fig. 3). There are few changes between the sensitized and hyperinsulinemic adipocytes in their basal state (Fig. 3b), but the 50 significantly differentially expressed genes are enriched in genesets annotated by DAVID(*18*) as primarily restricted to lipid and metabolism-related processes (Supplementary table 2). However, as opposed to the basal state, insulin-responsive transcriptional changes did display numerous differences between the sensitive and resistant adipocytes (Fig. 3b, 3c). The top differentially regulated genes after insulin stimulation in the sensitized adipocytes were largely also regulated in the resistant adipocytes, albeit with a diminished fold change (Fig. 3d).

**Fig. 3:**
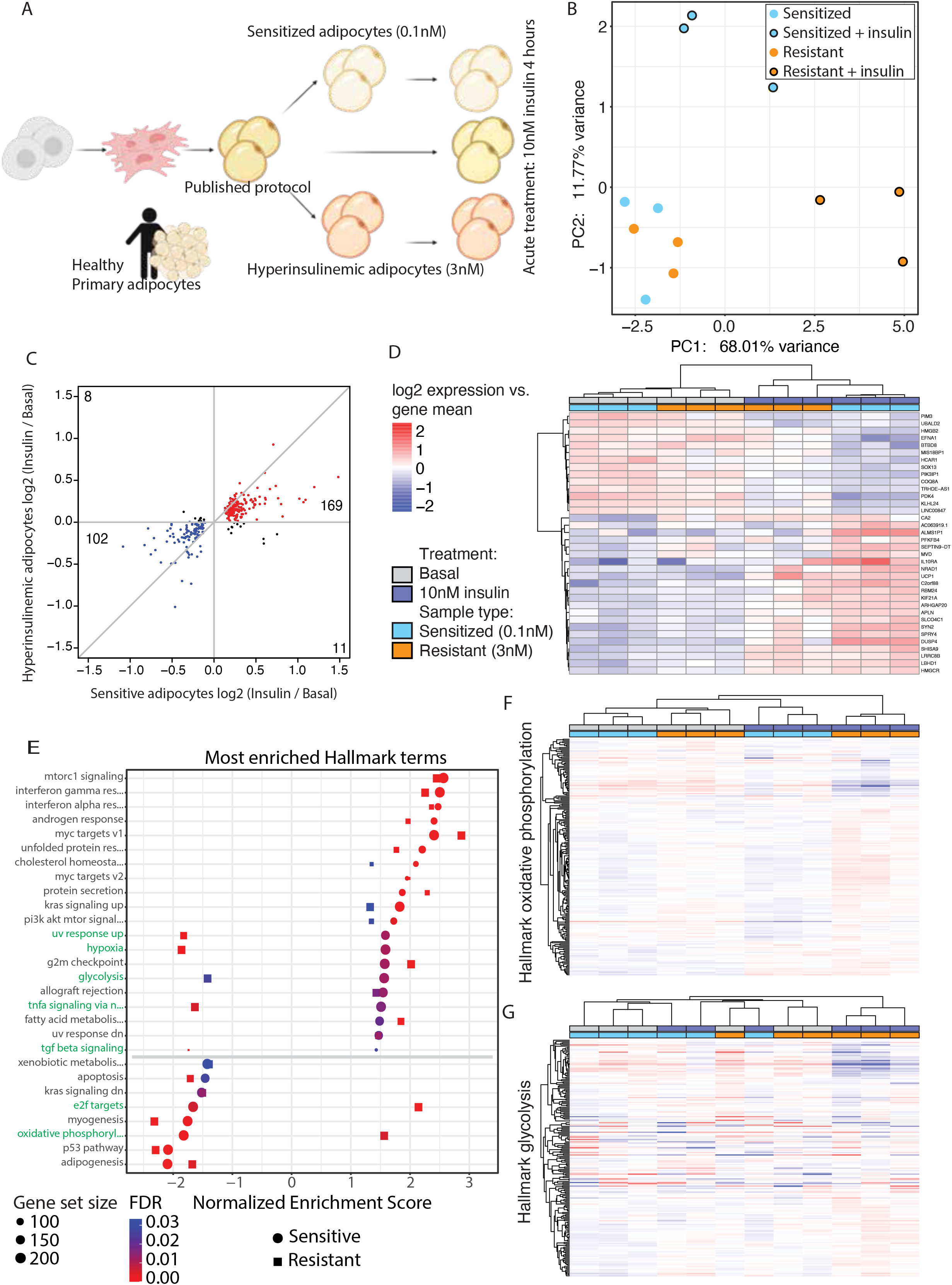
Induction of insulin resistance perturbs insulin-stimulated transcription A, Schematic indicating the experimental setup for the transcriptomic experiments. B, PCA plot showing the sensitized and resistant adipocyte in their basal or insulin-stimulated state. C, Scatterplot comparing fold change in sensitive and resistant adipocytes upon insulin stimulation, filtered on FDR<=0.05 in the sensitive adipocytes. Genes with a positive or negative fold change in both comparisons are colored red and blue respectively, genes with opposite fold changes appear in black. D, Heatmap of the genes most changed upon insulin stimulation, filtered by FDR<=0.05 and log2FC>0.5 in the sensitive adipocytes. E, GSEA results of Hallmark terms that are enriched upon insulin stimulation in both sensitive and resistant adipocytes. Ranked by NES score in the sensitive adipocytes. F, Heatmap of expression of genes in the oxidative phosphorylation Hallmark gene set. G, Heatmap of expression of genes in the glycolysis Hallmark gene set. Panel D, F, G use the same legend.

Given the lack of clear differences between the sensitized and insulin-resistant adipocytes in the basal state, we performed gene set enrichment analysis (GSEA)(*19*) on insulin-responsive genes in both of these conditions. We observed a large overlap in the most enriched Hallmark terms (Fig. 3e, Supplementary fig 3). As expected, many of the top enriched gene sets affected by insulin were related to metabolism (mTORC1 signaling, cholesterol homeostasis, PI3K AKT MTOR signaling, glycolysis, fatty acid metabolism, oxidative phosphorylation). Curiously, several gene sets were enriched in the opposite direction between sensitized and resistant adipocytes, with the most notable examples being the “glycolysis” and “oxidative phosphorylation” gene sets (Fig. 3e). In the sensitized adipocytes, we observed the expected metabolic response to insulin exposure; the expression of genes involved in glycolysis increased while oxidative phosphorylation related gene expression was decreased. In contrast, the resistant adipocytes exhibited the opposite response to insulin exposure — expression of glycolysis genes was decreased and oxidative phosphorylation genes increased with insulin — indicating an divergent regulation of metabolism. Gene expression changes in the oxidative phosphorylation gene set were generally mirrored between sensitized and resistant adipocytes, although there were several genes that were increased in the basal insulin-resistant state, which aberrantly further increased with insulin stimulation (Fig. 3f). Meanwhile, the resistant adipocytes in their basal state most closely resembled insulin-stimulated sensitive adipocytes with respect to glycolysis-related genes, while insulin stimulation in the resistant adipocytes wholly dysregulates the glycolytic transcriptional program (Fig. 3g). These results indicate that the sensitization protocol described here produced adipocytes similar in identity to published protocols with respect to primary tissue, but that our approach bolstered the transcriptional response to insulin. As expected, the most potentiated transcriptional pathways were related to metabolism, and these were adversely affected in the resistant adipocytes compared to the sensitive protocol.

## Discussion

The protocol developed here results in insulin-sensitive human adipocytes. By optimizing the timing and medium composition through a statistically driven Design-of-Experiments approach, we developed a simple, easily adoptable media formulation for producing adipocytes from hPSCs that show exquisite sensitivity to insulin signaling. Moreover, we have shown that insulin sensitivity can be dampened by exposing these cells to hyperinsulinemic conditions during the sensitization period, thus providing a new and physiologically relevant model of insulin resistance in human adipocytes. Additionally, our induction of insulin resistance did not produce extensive basal changes in transcription but did lead to a diverging response in gene expression upon insulin stimulation. This finding may suggest this model resembles progression towards insulin resistance in adipose tissue during hyperinsulinemia in humans. It also informs future research, as the lack of transcriptional changes between sensitive and resistant adipocytes strongly implies that the insulin resistance we observe in signaling and function is mediated at a protein or metabolic level. Additionally, these observed insulin-stimulated transcriptional differences in sensitive and resistant adipocytes are primarily in metabolic pathways, hinting that metabolism-related protein post-translation modifications such as oxidation or lipidation could be studied in our model system. Ultimately, this is the first human stem cell-derived adipocyte model that is insulin-responsive at physiologically relevant insulin concentrations and thus better mimics in vivo conditions. This unlocks a plethora of future research possibilities with all the advantages of human stem cell disease models.

## Materials and methods

### hPSC maintenance

hPSCs were maintained feeder-free on Matrigel (Corning 354234) in StemFlex medium (Thermo Fisher Scientific A3349401) and passaged as clumps using Versene solution (Thermo Fisher Scientific 15040066).

| hPSC identified | Cell line name | Source |
| --- | --- | --- |
| ESC#1 | Hues9 | HSCI (Harvard Stem Cell Institute) iPS Core Facility, Harvard University |
| ESC#2 | WA09 / H9 | WiCell |
| iPSC#1 | DiPS 1016 SeVA | HSCI (Harvard Stem Cell Institute) iPS Core Facility, Harvard University |
| iPSC#2 | WIBR599 | Whitehead Institute for Biomedical Research |

### Adipocyte differentiation

Differentiation of hPSCs into adipocytes was performed as published (*12*). Briefly, hPSCs were passaged and grown in suspension culture to form embryoid bodies. After one week, the embryoid bodies were plated on tissue culture-treated plastic and grown in DMEM + 10% fetal bovine serum. The outgrowing mesenchymal progenitor cells (MPCs) were used from passage 4 through 8. For adipocyte differentiation MPCs were infected with previously described lentivirus to express the transactivator rtTA and inducible expression of PPARg2. MPCs were passaged and grown to confluence, and then exposed to 700ng/mL doxycycline for two weeks in previously established A2 medium, and then one week without doxycycline in A2 medium. Afterward, adipocytes were exposed to the experimental treatment media for 5 days unless otherwise indicated.

### Design-of-Experiments

Media for the DoE experiments were prepared by using no-glucose DMEM (Thermo Fisher Scientific 11966025) and adding the components outlined in Supplementary Table 1. The selection of these formulations was made using JMP software (SAS) to maximize the experimental space created by the 4 factors. The design was a rotatable surface around a center point for the four factors. Factor levels were selected as +/-1 step sizes with log bases 10, 2, 5, and 2 for insulin, glucose, IGF-1, and albumin, respectively (Supplementary Table 1). Adipocytes were fed every other day with these media for 7 days before assays were performed. For modeling, the results of duplicate treatments for each condition were entered into JMP software). We performed least-squares fitting of linear dependencies for each factor and squared dependencies via self- and inter-factor crosses to determine any interacting effects on both glucose uptake and AKT2 phosphorylation, normalized to total protein for each condition. The model dependence for each of these linear and secondary factor interactions for both glucose uptake and AKT2 phosphorylation were reported as the parameter coefficient, and the significance of each contribution to the model was reported as the t-ratio p-value.

### Insulin stimulation assays

All experiments were performed with a no-glucose DMEM as the assay medium (Thermo Fisher Scientific A1443001). At the endpoint of differentiation and medium treatment, adipocytes were rinsed with assay medium, and then washed for 10 minutes at 37C. For ELISAs, adipocytes were treated with assay medium +/-insulin for 10 minutes and harvested in cell lysis buffer (Cell Signaling Technology #9803) with phosphatase inhibitor (Thermo Fisher Scientific 78442). For glucose uptake assays, adipocytes were treated with assay medium +/-insulin for 40 minutes and 2μM 2-deoxy-glucose and processed as per manufacturer’s instructions.

### AKT2 ELISA

After cell lysis, protein samples were quantified and diluted as appropriate for the ELISA (∼100-200ng/mL protein). PathScan® Phospho-Akt2 (Ser474) and Total Akt2 Sandwich ELISA kits were used to quantify AKT2 levels (Cell Signaling Technology #7048 and #7046 respectively) as per manufacturer’s instructions, by colorimetric reading at 450nm on a Thermo Fisher Multiskan Go plate reader. Results are plotted as the OD450 ratio of phosphorylated over total AKT2, or as fold change compared to the basal unstimulated state.

### Glucose uptake

Glucose uptake was measured using the Glucose Uptake-Glo™ Assay (Promega J1342) per the manufacturer’s instructions. Briefly, after assay medium treatment, 0.5 volume equivalents of lysis buffer was added, followed by 0.5 volume neutralization buffer and 1 volume of the prepared 2DG6P detection reagent. Luminescence was measured to quantify the amount of 2DG taken up. Results are normalized by total protein per well and plotted as relative luminescence units (RLU) or as fold change compared to the basal unstimulated state.

### Protein quantification

Total protein was quantified for all samples using the Pierce™ BCA Protein Assay Kit (Thermo Fisher Scientific 23225) as per the manufacturer’s instructions.

### Immunofluorescence and TIRF

Differentiated adipocytes were fixed with 10% neutral buffered formalin for 15 min and then washed twice with PBS. Cells were permeabilized for 10 min with 0.1% Triton X-100 and incubated in blocking solution consisting of 2% bovine serum albumin (BSA) in PBS for 1 hr at room temperature. After blocking, the cells were incubated in primary antibodies (anti-Glucose Transporter GLUT4 (Abcam ab654 1:500 dilution) or rabbit anti-C/EBPα (Cell signaling technologies D56F10 1:200 dilution) for 1 hr at room temperature in 0.2% BSA and 0.1% Triton X-100 in PBS. After primary incubation, samples were washed three times for 5 min each in 0.05% Tween-20 in PBS. After washing, cells were incubated in secondary antibodies (donkey anti-rabbit 488 (Life Technologies A-21206 1:500)) for 30 min at room temperature in 0.2% BSA and 0.1% Triton X-100 in PBS. In addition, DAPI (Life Technologies A-62248 1:500) and HCS LipidTOX™ Deep Red Neutral Lipid Stain (ThermoFisher Scientific H34477 1:500) were added to the secondary stain solution for counterstains of the nuclei and lipid droplets, respectively. After secondary and counterstain incubation, samples were washed three times for 5 min each in 0.05% Tween-20 in PBS. GLUT4 total internal reflection (TIRF) imaging was performed on a Nikon Ti-E inverted microscope with a Yokogawa CSU-X1 spinning disk confocal scan head, Andor iXon 897E EM-CCD camera, and Andor FRAPPA TIRF photomanipulation system using a 60X plan apo TIRF objective. C/EBPα epifluorescence imaging was performed on a Nikon TE2000 inverted microscope with a Hamamatsu Orca-ER CCD camera using a 10X plan fluor objective. For GLUT4 TIRF imaging quantification, single adipocytes in the stimulated and unstimulated condition were selected at random using the lipid droplet and then centered around the nuclei. Approximately 30 individual cells were selected and imaged between 2 biological replicates. The AlexaFluor-488 GLUT4 area was outlined in ImageJ (NIH) and the integrated intensity was measured and plotted. For C/EBPα imaging and quantification, 3 representative areas were selected from three replicate differentiations using the DAPI counterstain. The differentiation efficacy for each cell line in using the previously published medium and the sensitization medium is was reported as C/EBPα per unit area by average the 3 representative areas together for a single n and generating n=3 for the replicate differentiations.

### Primary adipocytes

Primary adipocyte RNA was procured from Zenbio. The following samples were used:

| Catalog # | Lot # | Description |
| --- | --- | --- |
| RNA-A10-1 | LM082310A | Human culture adipocyte RNA <24.9BMI |
| RNA-A10-2 | L071498 | Human culture adipocyte RNA 25-29.9BMI |
| RNA-A10-2 | L090198 | Human culture adipocyte RNA 25-29.9BMI |

### RNA sequencing

We mapped the reads to the hg38 version of the human genome, containing only canonical chromosomes, with STAR(*20*) using a GTF file downloaded from ESEMBL version GRCh38.99. We set the overhang parameter to 50 and the “alignIntronMax” parameter to 95000. We assigned reads to genes with featureCounts(*21*) with parameters “-p -s 2” and the same GTF file used on the mapping step. We normalized the counts and performed principal components analysis (PCA) with DESeq2(*22*) using “vst” on the variance stabilization step. Samples were contrasted in PCA space by Euclidean distance. We performed differential expression (DE) analysis with DESeq2 without log fold change shrinkage. We used the statistic from the DE analysis to rank the genes and run the pre-ranked gene set enrichment analysis (GSEA)(*19*), with the hallmark gene sets version 7.4. We used Cluster 3.0(*23*) to cluster the genes and samples displayed on the heatmaps.

## Acknowledgements

This study was supported by Novo Nordisk to M.F. and R.J., the National Institute of Health to R.J. (1U19AI131135-01; 5R01MH104610-21) as well as the National Insitute of Biomedical Imaging and Bioengineering at the National Institute of Health (T32 EB016652) to A.S.K. and D.J.M. We thank the Whitehead Institute genome technology core for the RNA-sequencing. Some of the figures were made with the help of biorender.com.

## Author contributions

MF, ASK, JFJ and RJ designed the study. MF and ASK performed experiments. MF, ASK, and MIB analyzed data. All authors contributed to the manuscript.

## Figure legends

**Supplemental fig. 1:**
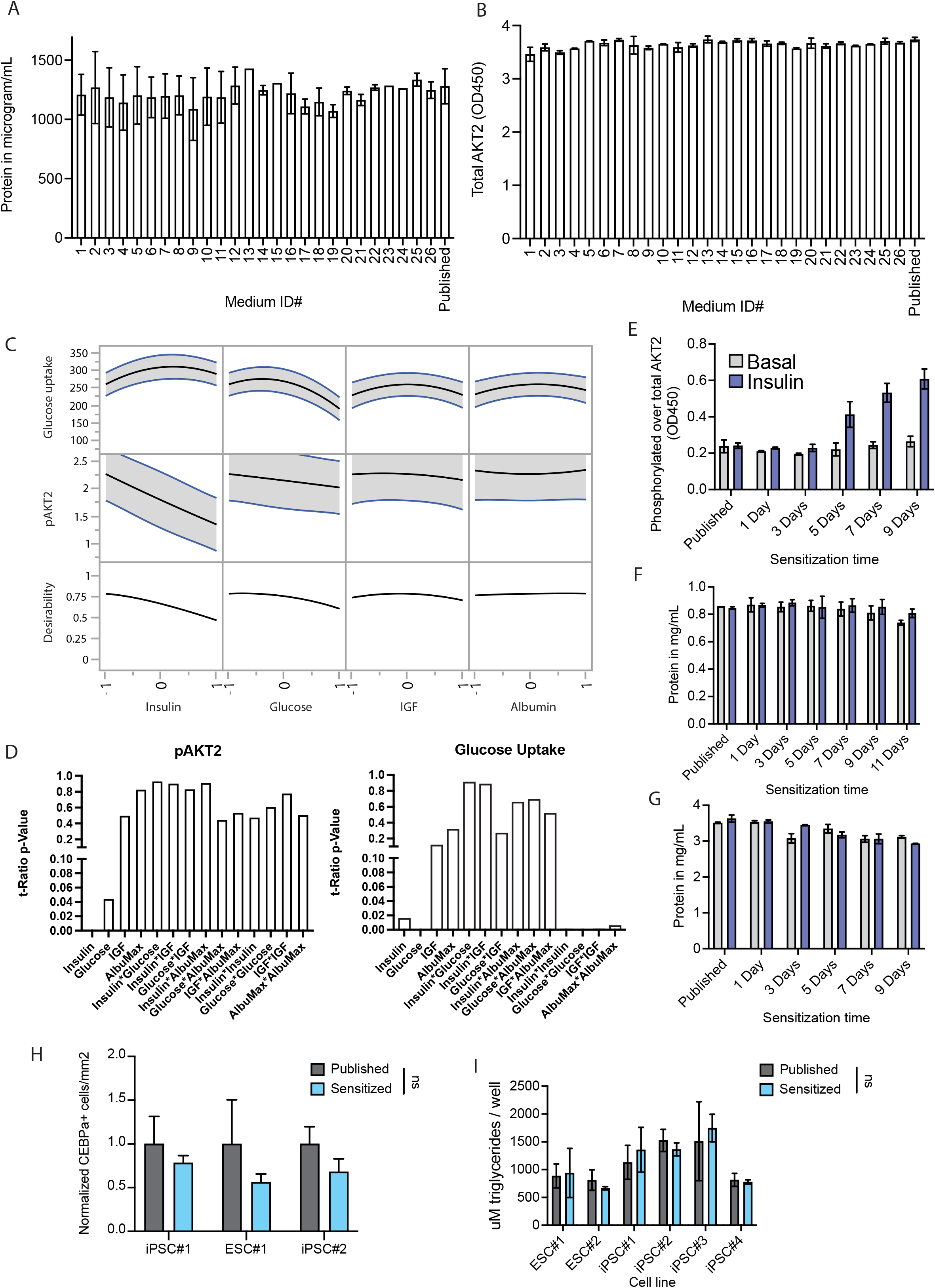
A, Total protein content of samples used in the DoE screen and the published protocol medium. B, Total AKT2 ELISA measurement of samples in the DoE screen. C, Profile curves for each factor illustrating the influence on glucose uptake and phosphorylation of AKT2 throughout the experimental space of the DoE. D, t-Ratio p-values indicating the significance of each factor in the DoE screen for phosphorylated AKT2 (left) and glucose uptake (right). E, Timecourse of sensitization with the DoE-optimized media measuring phosphorylation of AKT2 at baseline and after insulin stimulation. Results are normalized to total AKT2 for each sample. F, Total protein content of each sample during the glucose uptake timecourse (Fig. 1f). G, Total protein content of each sample during the AKT2 timecourse (Supp. Fig. 1e). H, Quantification of adipocyte marker CEBPA-positive nuclei per area in published and sensitized-protocol adipocytes. I, Triglyceride content per sample of published and sensitized adipocytes in several cell lines. All bar graphs depict the mean with error bars representing s.d., n=2 biological replicates for A-G, n=3 for H,I.

**Supplemental fig. 2:**
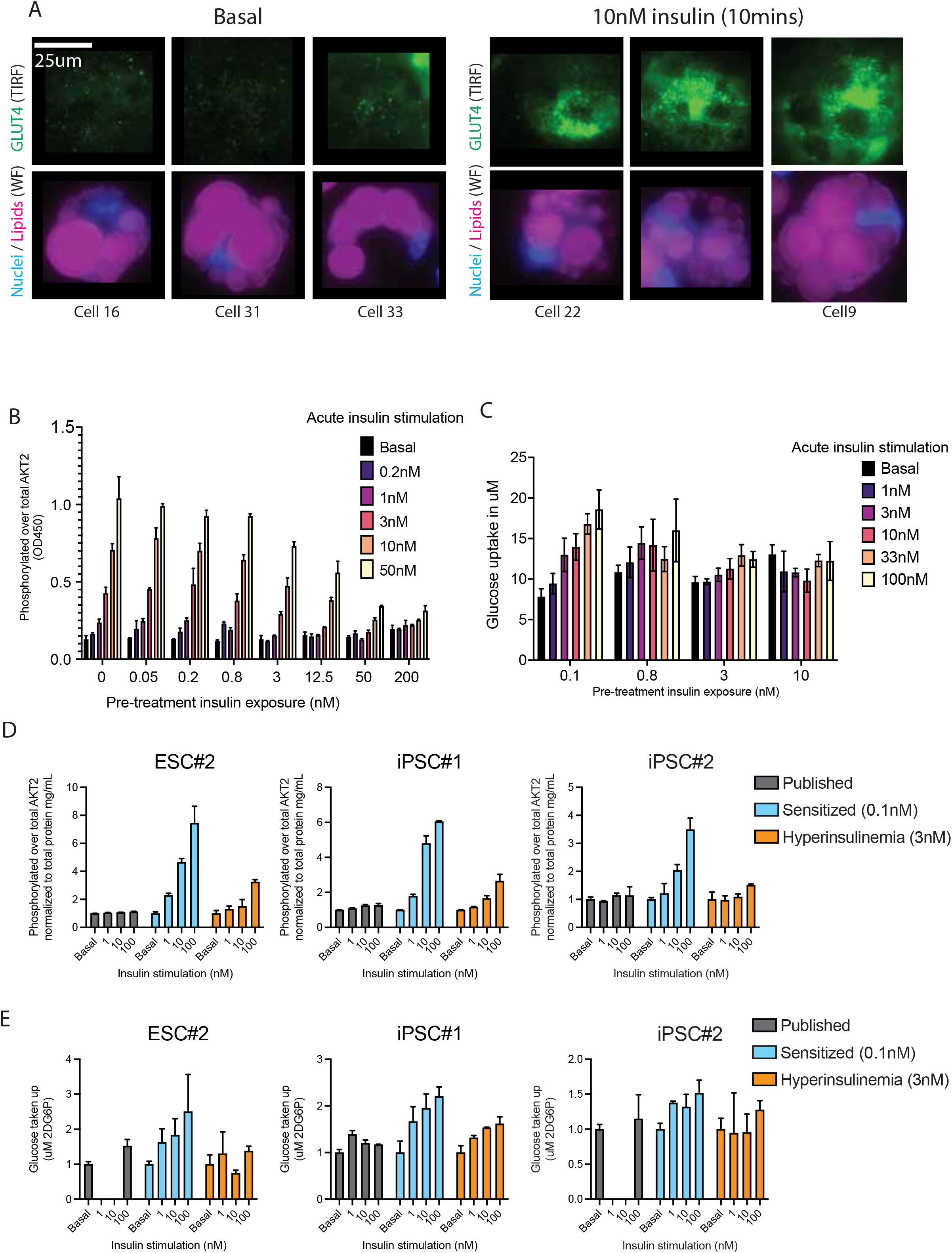
A, Representive images of the TIRF quantification in fig. 1d. For each condition, the three cells closest to the mean are shown. B, Full panel of pre-treatment insulin exposure concentrations and acute stimulation dose-response measuring phosphorylated AKT2 normalized to total AKT2, related to figure 2f. C, As in B but measuring glucose uptake normalized to total protein, related to figure 2g. D, Phosphorylated AKT2 measurements normalized to total AKT2 for 3 additional hPSC lines. E, As in D but measuring glucose uptake normalized to total protein. All bar graphs depict the mean with error bars representing s.d., n=2 biological replicates for B,C, n=3 for D,E.

**Supplemental fig. 3:**
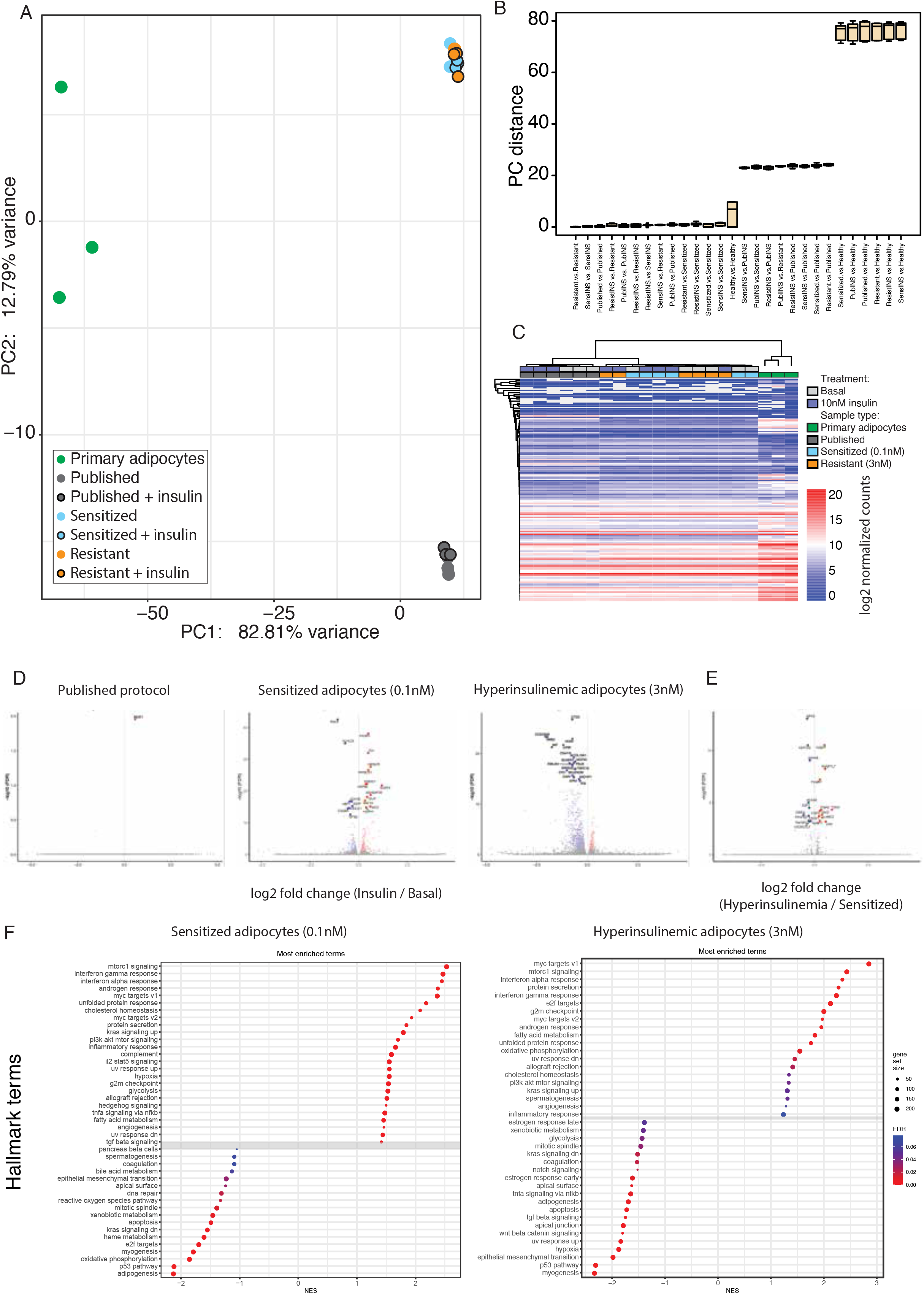
A, PCA plot of all samples assessed with RNA-sequencing. B, Measurement of PCA distances between all sample groups. C, Total expression heatmap of adipocyte genes identified through the Human Protein Atlas. D, Volcano plots of published, sensitized, and hyperinsulinemic adipocytes showing log2 fold change of genes upon insulin stimulation. Genes with FDR<=0.05 are colored, and the top 20 significant genes are labeled. E, Volcano plot of gene changes between sensitized and hyperinsulinemic adipocytes. Genes with FDR<=0.05 are colored, and the top 20 significant genes are labeled. F, GSEA results of all Hallmark terms that are enriched upon insulin stimulation in the sensitive (left) and resistant (right adipocytes, ranked by NES score, related to figure 3e.

---

Supplemental table 1: Design of Experiments variable inputs

Supplemental table 2: Results of DAVID clustering on the significant genes between adipocytes from the published and sensitive protocol.

